# RBM15-MKL1 fusion protein promotes leukemia via m6A methylation and WNT pathway activation

**DOI:** 10.1101/2025.02.19.638991

**Authors:** Madeline Y. Mayday, Giulia Biancon, Manyi Wei, Christian Ramirez, Irene Moratti, Andreas P. Pintado-Urbanc, Lin Wang, Jether Amos Espinosa, Matthew D. Simon, Yaara Ofir-Rosenfeld, Oliver Rausch, Toma Tebaldi, Stephanie Halene, Diane S. Krause

## Abstract

Acute megakaryoblastic leukemia driven by the RBM15-MKL1 fusion protein (RM-AMKL) is the only known recurrent mutation involving the N6-methyladenosine (m6A) writer complex. Dysregulation of m6A modification affects RNA fate and is linked to oncogenesis. Inhibition of m6A deposition via inhibition of the METTL3 writer protein has anti-tumour activity, but the mechanism underlying its efficacy and cancer specificity remains unclear. We treated murine RM-AMKL cells with a novel METTL3 inhibitor, STM3675, and showed apoptosis *in vitro* and prolonged survival of mice transplanted with RM-AMKL, implicating m6A as an essential component of AMKL and identifying Wnt signalling as a key driver of leukemogenesis. To elucidate the mechanism by which m6A contributes to leukemogenesis we employed a multi-omic approach, combining transcriptome-wide assessment of RNA binding, methylation and turnover. We show for the first time that RM retains the RNA-binding and m6A-modifiying functions of its RBM15 component, while also selectively regulating distinct mRNA targets, particularly genes involved in Wnt signalling including Frizzled. Frizzled genes are upregulated by RM and downregulated in RM-AMKL cells in response to METTL3 inhibition, providing an m6A-dependent explanation for their upregulation. Direct Frizzled knockdown reduced RM-AMKL growth, which was partially rescued by treatment with a β-catenin agonist, underscoring a functional role of Wnt signalling in RM-AMKL. Human AMKLs show elevated Wnt pathway and Frizzled gene expression, highlighting the relevance of our work. Together, our findings reveal that RM-specific m6A modifications and activation of Wnt signalling are critical drivers of RM-AMKL, highlighting these pathways as potential therapeutic targets.

**Key Points:**

- RM retains functional abilities of RBM15 and additionally interacts with Wnt-related transcripts to increase expression of Fzd proteins.
- The METTL3 writer complex and WNT signalling pathways are essential for RM-driven leukemia.

## Introduction

Acute megakaryoblastic leukemia (AMKL) is a subtype of acute myeloid leukemia (AML) characterized by failed megakaryocyte maturation and expansion of abnormal megakaryoblasts. AMKL primarily affects children, accounting for 4-15% of pediatric AML cases, and although survival rates for many pediatric cancers have improved, survival for neonates with AMKL remains variable^1^. Current treatments are non-specific, highly toxic, and have limited efficacy. The dearth of data regarding the mechanisms of leukemogenesis represents an obstacle to improving therapies for AMKL.

AMKL with the recurrent t(1;22) translocation, which encodes the RBM15-MKL1 (RM) fusion protein ^2,3^, provides novel insights into leukemogenesis. RBM15 (RNA-binding motif protein 15) plays a critical role in RNA N6-methyladenosine (m6A) modification^4–8^ by recruiting the m6A writer complex, which has catalytically active methyltransferase 3 (METTL3) at its core. Recruitment by RBM15 is essential for establishing the m6A epitranscriptome, influencing RNA fate by altering transcript stability^5,9^. In RM, RBM15 is fused to MKL1 (megakaryoblastic leukemia 1), which functions as a transcriptional co-activator of serum response factor (SRF) to regulate genes related to cell cycle and megakaryocyte differentiation^10–15^.

RM retains the functional domains of both RBM15 and MKL1, and as such it may direct leukemogenesis via interactions with the genome, transcriptome, and epitranscriptome. To date, RM has been linked to dysregulation of SRF target genes^16^, interactions with histone H3K4 methyltransferase Setd1b^17^, and dysregulation of Notch signalling^18^, though this has been challenged by data from primary patient samples^19^. Thus, the driving mechanisms behind RM-AMKL remain unclear.

Given RBM15’s role in m6A modification, we investigated whether targeting the epitranscriptome could provide mechanistic rationale for novel targeted approaches. Modulation of the m6 epitranscriptome via inhibition of METTL3 has been investigated as a therapy for cancer, and potent, highly specific METTL3 inhibitors have shown promise in AML^20^. Since RM is the only recurrent disease-related genetic alteration directly involving a member of the m6A writer complex, it serves as an important model system for investigation of fundamental biological mechanisms that are also relevant to more prevalent diseases.

We show that METTL3 inhibition promoted apoptosis and differentiation of murine RM- AMKL cell lines and prolonged survival of AMKL-engrafted mice. To understand the mechanistic basis for this response, we leveraged high-resolution multi-omic analyses to examine how RM influences RNA and m6A modification. We integrated anti-RM and anti- RBM15 with anti-m6A RNA crosslinking and immunoprecipitation and show for the first time that RM retains the ability to bind RNA and to direct target-specific m6A modification. RNA-sequencing and TimeLapse-sequencing reveal that RM-specific RNA targets are differentially expressed and exhibit altered stability. Our integrative analysis suggests that RM affects *Frizzled* gene methylation, expression, and protein levels as well as cell survival, and that these alterations are reversed by METTL3 inhibition, linking RM directly to Wnt signalling and nominating METTL3 inhibition as a RM-targeted therapeutic opportunity.

Our studies provide the first deep mechanistic understanding of RM-AMKL and its interactions with RNA, opening new avenues of investigation toward development of targeted treatments for AMKL.

## Methods

### Cell lines

Human erythroid leukemia (HEL) cells were transfected as described^14^ to produce control cells and cell lines with inducible RM-FLAG and RBM15-FLAG. 6133 cells^18^ were a gift of Thomas Mercher and Gary Gilliland. The CAOM cell line was generated by us from a Rbm15-MKL1 knockin mouse^18^ that spontaneously developed leukemia.

### Mouse models

400,000 CAOM cells were transplanted into sub-lethally irradiated (600cGy) mice. Recipient mouse blood was analyzed to validate leukemia engraftment. Animals were euthanized when moribund and analyzed for engraftment in bone marrow and spleen.

### METTL3 inhibitor

Characterization of STM3675 was performed as described^20,21^. For *in vitro* use, STM3006 and STM3675 were diluted to achieve final concentrations as indicated. For *in vivo* use, 10mg/ml STM3675 stock solution was administered at 100mg/kg via oral gavage.

#### Cell toxicity assay and morphologic analysis

Cells were treated with Vehicle or serial concentrations of STM3006 or STM3675 and measured using CellTiter-Glo®. The IC50 was calculated by nonlinear regression. Cells treated with 1µM were assessed after Wright-Giemsa staining.

### Flow cytometry

HEL cell ploidy was determined using Propidium Iodide as described^14^ and cell maturation was assessed using CD49b-PE and CD235A-APC. 6133 and CAOM apoptosis was assessed with the Annexin V Apoptosis Detection Kit. Differentiation was assessed using 7-AAD, anti-ckit-APC-Cy7, anti-CD41-FITC or anti-CD61-FITC. Murine bone marrow progenitors were sorted as previously described^22,23^. Megakaryocyte progenitors were identified as Lin^−^cKit^+^Sca^−^CD150^+^CD41^+^ and erythroid progenitors as Lin^−^cKit^+^Sca^−^CD150^+^FcgRI/II^−^CD105^+^.

### Sequencing experiments and analyses

All libraries underwent 100bp paired-end sequencing on the Illumina NovaSeq 6000 at the Yale Center for Genome Analysis.

#### RNA-seq

For HEL and progenitor cells, RNA was isolated using the RNeasy Mini Kit. For 6133 and CAOM cells, RNA was isolated using Trizol. RNA underwent poly-A selection or rRNA depletion and reverse transcription and library preparation using the KAPA mRNA HyperPrep kit.

After quality control and filtering, reads were aligned to GRCm39 or GRCh38 with STAR using the Gencode M31 and 37 gene annotation, respectively. edgeR was used for normalization (TMM method) and identification of differentially expressed genes (absolute log2 fold change>0.75, FDR<0.05).

#### Enhanced crosslinking and immunoprecipitation sequencing (eCLIP-seq)

Anti-FLAG eCLIP-seq was performed according to published protocols^24–26^ with minor adaptations (See Supplemental Methods). For m6A eCLIP-seq, mRNA was isolated, fragmented, incubated with ribonuclease inhibitor and anti-m6A antibody, and crosslinked. Antibody-RNA complexes were immunoprecipitated and eCLIP performed as above.

After quality filtering, peak calling was performed with PureCLIP^27^. Read counts for each peak region were normalized and differential analysis was performed as for RNA-seq using RBM15 regions as controls. Differentially bound or methylated regions were identified as p<0.05.

To identify “couplets”, each binding site was paired with the nearest m6A site. Differential couplets were identified as p<0.05 for anti-FLAG or anti-m6A eCLIP. Delta delta couplets were identified among differential couplets by scaling RM vs RBM15 fold-changes on RNA-fold changes and selecting couplets with a resulting scaled fold-change ≥ 1 or ≤ -1 in anti-FLAG or anti-m6A eCLIP.

#### TimeLapse-seq

Experiments were performed as described^28^. HEL cells were supplemented with 100µM 4-thiouridine during the last 2h of culture. Reads were processed as described^28^. bakR and DESeq2 were used to calculate differences in kinetic parameters and gene expression levels^29,30^. Transcripts with P_adj_<0.05 in k_deg_ and/or k_syn_ across conditions were called as differentially stable and/or synthesized, respectively.

#### AMKL and AML patient samples

RNA-Seq gene counts from AMKL, AML, and healthy control samples were obtained^1,31,32^ and processed as for RNA-seq above. Wnt pathway activation scores for each sample were obtained from the cellmarkeraccordion package^33^.

## Results

### Novel METTL3 inhibitor is effective against mouse models of RM-AMKL

Pharmacologic inhibition of METTL3 induces differentiation and apoptosis of AML cells by limiting translation of oncogenes^20^. In solid tumor models, METTL3 inhibition induces a cancer cell-intrinsic innate immune response^21^. Given the role of RBM15 in the m6A writer complex, we tested whether pharmacological inhibition of METTL3 affected RM- AMKL. We used two METTL3 inhibitors (**Figure 1A**); STM3006 is a highly potent and specific METTL3 inhibitor that blocks m6A modification at nanomolar concentrations but is rapidly metabolized *in vivo*^21^. We therefore employed a previously undisclosed METTL3 inhibitor, STM3675, that shows equally high specificity and potency against METTL3 but has good bioavailability *in vivo*. STM3675 (**Figure 1A**) inhibits recombinant METTL3 activity with an IC50 of <6nmol/L, a value restricted by the lower detection limit (**Figure S1A**). Surface plasmon resonance revealed STM3675 binding to METTL3 with a Kd of 22pmol/L (**Figure S1B**). STM3675 reduced m6A modification of polyA-RNA in cells with an IC50 of 4.5nmol/L (**Figure 1B**). Like STM3006^21^, STM3675 shows greater than 1,000- fold selectivity for METTL3 compared to other RNA methyltransferases (**Figure S1C**). Pharmacokinetics demonstrated high oral bioavailability and metabolic stability in mice (**Figure S1D**). Analysis of m6A levels on polyA-RNA *in vivo* showed maximal inhibition at 10-100mg/kg (**Figure S1E**).

**Figure 1.**
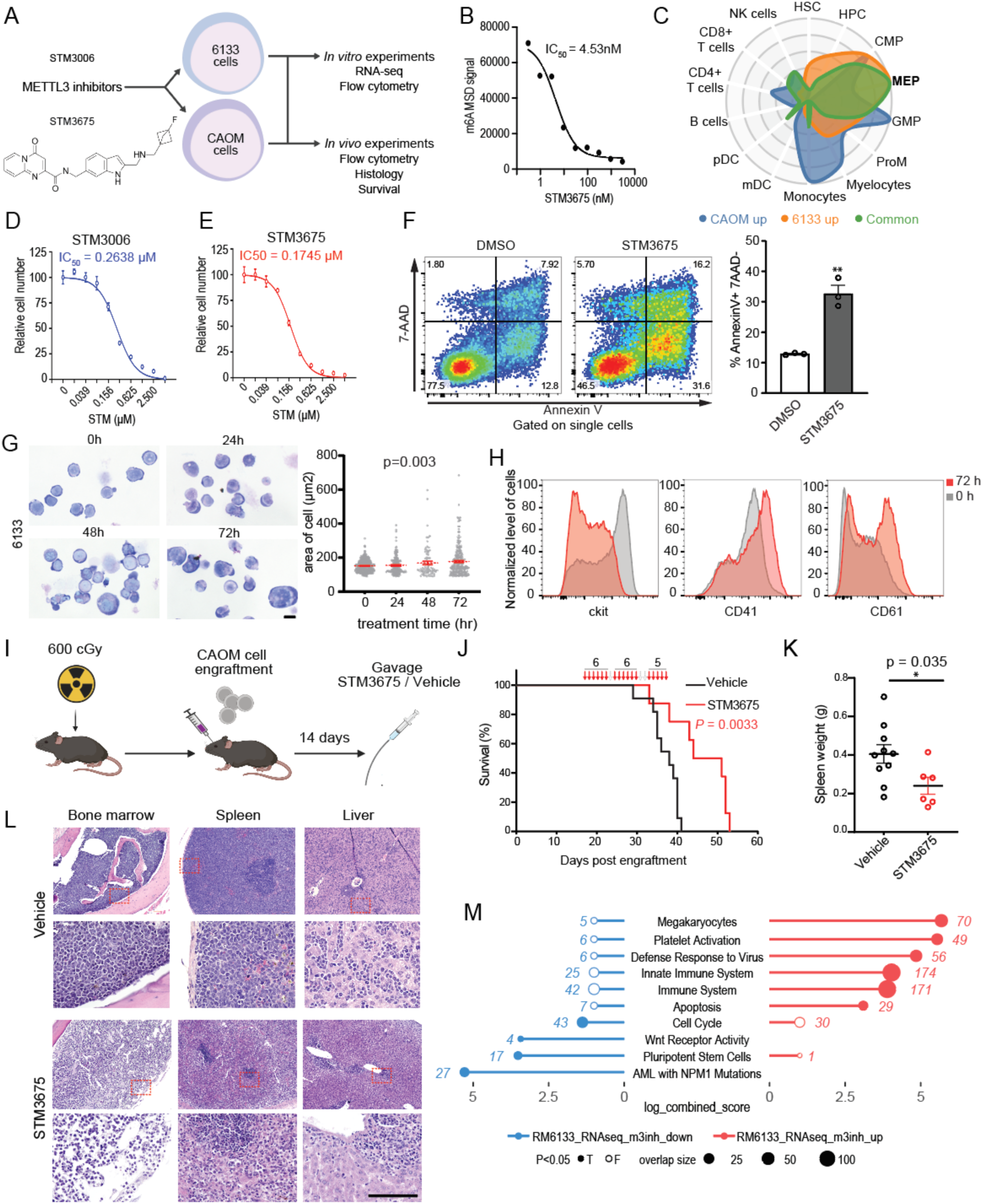
Inhibition of Mettl3 induces AMKL blast death and prolongs survival in leukemic mice. (a) Schematic of experiments and structural formula of STM3675. (b) m6A quantification and IC50 determination as measured by m6A electroluminescence (ECL) ELISA. (c) 6133 and CAOM cells have a shared megakaryoblastic expression profile as shown via CellRadar analysis (https://karlssong.github.io/cellradar/), comparing genes differentially upregulated in CAOM (blue) or 6133 (orange) or commonly expressed (green) to normal human hematopoiesis datasets from Bloodspot. (d,e) STM3006 and STM3675 are toxic to 6133 cells in vitro. Total cell number was normalized to vehicle. IC50 was calculated by fitting toxicity curve using a nonlinear method. n = 3 biological replicates. (f) Flow-cytometric assessment and quantification of apoptosis (Annexin V + 7AAD -) in response to vehicle or STM3675 (1µM). A representative experiment of three is shown, performed in triplicates. Data are represented as mean ± SEM ; the p-values were calculated using independent two-tailed Student’s t test. **p < 0.01. (g) Wright-Giemsa staining and quantification of cell area (µm^2^) of CAOM cells treated with Vehicle (0 hour) or STM3675 for 24, 48, 72 hours. Scale bar, 10 μm. p-values calculated with one-way ANOVA with Welch’s test, p = 0.003. (h) Flow-cytometric assessment and quantification of c-kit, CD41 and CD61 expression in 6133 cells treated with vehicle (0 hour) or STM3675 (1µM) for 24, 48 or 72 hours. A representative experiment of three is shown. (i) In vivo treatment of CAOM-engrafted, sublethally irradiated Pep3B recipient mice with vehicle or STM3675 via oral gavage. (j) Survival curve of mice engrafted with CAOM cells treated with vehicle (black) or STM3675 (red). *** P = 0.0033. (k) Spleen weights of vehicle versus STM3675 treated animals at time of censorship. *P<0.05. (l) Representative histologic images of bone marrow and spleen are shown from a vehicle and STM3675 treated animals after completion of treatment at time of censorship. Scale bar represents 100 μm. Quadrilateral frame labels the amplification area showed below. (m) Functional enrichment analysis of differentially expressed genes from 6133 cells treated with STM3675 or vehicle (1,950 upregulated genes and 1,015 downregulated genes). Data are represented as mean ± SEM and representative of at least three independent experiments; the p values were calculated using independent two-tailed Student’s t test (f,k) or one-way ANOVA with Welch’s test (g). n.s., not statistically significant; *p < 0.05 **p < 0.01; ***p < 0.001. *See also Figure S1*.

### Mettl3 inhibition stalls murine Rbm15-MKL1-driven AMKL *in vitro*

Since primary RM-AMKL patient samples are scarce, we used murine 6133 RM-AMKL cells^18^ to determine whether the m6A epitranscriptome represents a potential therapeutic target. We additionally generated a second murine RM-AMKL cell line, “CAOM”, from the same mouse model^18^ that developed spontaneously with a 17-month latency and that is serially transplantable. CAOM cells grow short-term *in vitro* in Scf-containing medium, have blast morphology, lack myeloid lineage markers (**Figure S1F**) and have low CD41 expression. 6133 and CAOM cells have distinct gene expression profiles but share a megakaryoblastic signature (https://karlssong.github.io/cellradar/)^34^ (**Figure 1C)**. These two murine RM-AMKLs mimic the heterogeneity encountered in AMKL patients^1^ and serve as physiologically relevant models for mechanistic investigations of RM-AMKL.

*In vitro* treatment with STM3006 and STM3675 for 5 days killed 6133 cells with IC50s of 0.264µM and 0.175µM, respectively (**Figures 1D** and **1E**). Given their equipotency, we focused further experiments on bioavailable STM3675. STM3675 induced apoptotic cell death (**Figure 1F**) and increased differentiation with downregulation of c-kit and upregulation of CD41 and CD61 (**Figures 1G, 1H** and **S1G**). STM3006 and STM3675 had similar effects on CAOM cells (**Figures S1H**, **S1I**, **S1J**, and **S1K**).

### Mettl3 inhibition prolongs survival in a syngeneic murine AMKL transplant model *in vivo*

We next assessed the *in vivo* efficacy of STM3675. 6133 cells grow as a Scf-dependent cell line^18^ but show poor *in vivo* engraftment. Therefore, we focused on CAOM cells which provide a reliable *in vivo* model. We transplanted CAOM cells into sub-lethally irradiated recipient mice and treated with Vehicle or 100mg/kg STM3675 via oral gavage over 21 days (**Figure 1I**). Vehicle-treated mice succumbed to disease within 41 days (median 38 days); mice treated with STM3675 showed prolonged survival to 52 days (median 51 days, **Figure 1J**). At sacrifice, spleens were significantly smaller with STM3675 treatment (**Figure 1K**). CAOM engraftment and treatment effects were validated via histology (**Figure 1L**). These data suggest that Mettl3-mediated m6A modification is essential for RM-AMKL.

### Mettl3 inhibition induces gene expression signatures related to differentiation and decreases oncogenic pathways including Wnt signalling

48-hour *in vitro* treatment with STM3675 (1µM) or Vehicle promoted significant gene expression changes in 6133 and CAOM cells. Upregulated genes in both models were enriched for mature megakaryocyte and platelet signatures, consistent with our flow data, as well as antiviral immune response pathways, known to be upregulated in response to Mettl3 inhibition^21,35,36^. Downregulated genes were enriched for stem cell and AMKL signatures as well as Wnt signalling (**Figure 1M** and **S1L**), suggesting that Mettl3- mediated m6A modifications affect the Wnt pathway.

### RBM15-MKL1 expression in HEL cells recapitulates AMKL differentiation block and activates RM-specific transcriptome

Because RBM15 is essential for m6A modifications, RM-AMKL offers a distinct opportunity to reveal the m6A-driven mechanisms promoting leukemogenesis. We therefore sought to identify how RM interacts with RNA differently from RBM15. Although 6133 and CAOM represent physiologically relevant models of AMKL, further mechanistic studies are hindered by the presence of both untagged fusion protein and wildtype Rbm15. Therefore, we generated HEL cells with doxycycline (dox)-inducible expression of FLAG-tagged RBM15 (HEL-RBM15) or FLAG-RM (HEL-RM) (**Figures 2A, 2B,** and **S2A**), allowing specific immunoprecipitation of each protein. HEL cells undergo megakaryocyte-like differentiation in response to 12-O-Tetradecanoylphorbol-13-acetate (TPA)^37,38^. We have previously shown that MKL1 over-expression (HEL-MKL1) promoted TPA-induced megakaryocytic differentiation. While dox-induced RBM15 did not have an effect (**Figure S2B**), expression of RM impaired megakaryocytic differentiation (**Figures 2C** and **2D**), a hallmark of AMKL.

**Figure 2.**
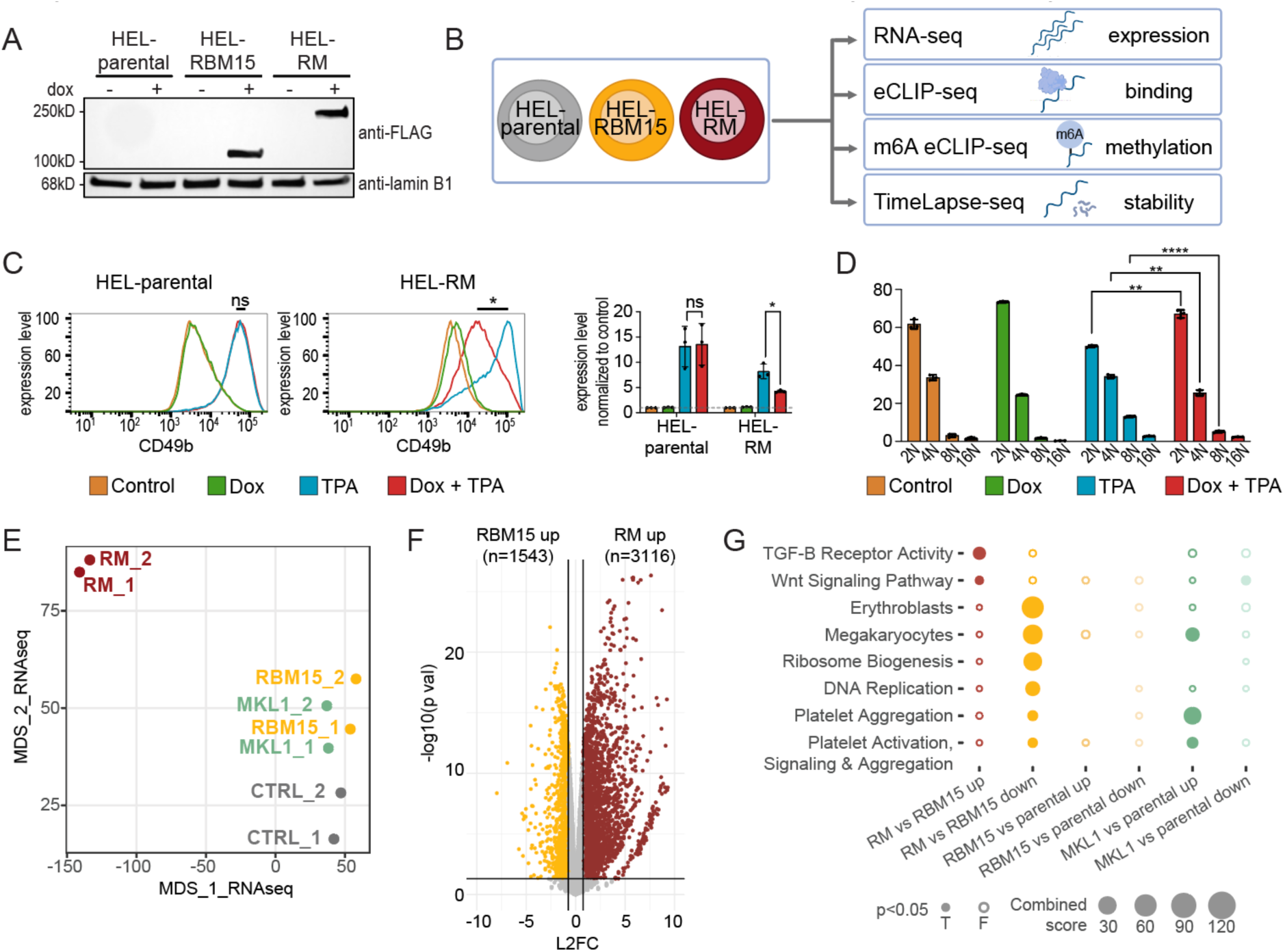
HEL cell model expressing RM recapitulates AMKL block in differentiation. (a) Western of nuclear lysate from HEL-RM, HEL-RBM15, and HEL-parental cells after 24h +/- dox, IB: anti-FLAG or lamin B1. (b) Schematic overview of sequencing experiments. (c) Flow cytometry on HEL-parental (left) and HEL-RM (right) without dox (orange), with dox (green), with TPA (blue), and with dox and TPA (red) shows decreased expression of megakaryocyte maturation marker CD49b in HEL-RM cells treated with dox and TPA (red) compared to TPA alone (blue). (d) Ploidy assays show reduced polyploidization (increased 2N and decreased 4N and 8N) in HEL-RM cells treated with dox and TPA (red) compared to TPA alone. (e) MDS plot of RNAseq from HEL-RM, HEL-RBM15, HEL-MKL1, and HEL-PCDNA6 cells. (f) Volcano plot of DEGs from HEL-RM (red, 3,116 DEGs) vs HEL-RBM15 (yellow, 1,543 DEGs). (g) Functional enrichment analysis of genes differentially regulated by RM vs RBM15, RBM15 vs parental, and MKL1 vs parental. Error bars represent mean ± SD, n=3, relevant p values are plotted (ns not significant, *p<0.05, **p<0.01, ****p<0.0001, 2-way ANOVA with Fisher’s LSD). *See also Figure S2*.

RNA sequencing on dox-induced HEL-RM, HEL-RBM15, HEL-MKL1, and HEL-parental cells revealed significant RM-specific transcriptional changes. HEL-RBM15 and HEL- MKL1 samples clustered with HEL-parental cells with small overall changes in gene expression (**Figure 2E**). RM expression separated HEL-RM cells from other conditions and confirmed compromised megakaryocyte differentiation (**Figures 2E**, **S2C, S2D** and **S2E**). Analysis of HEL-RM vs HEL-RBM15 identified similar gene expression changes to HEL-RM vs HEL-parental (**Figures 2F** and **S2F**). We therefore considered HEL-RBM15 an appropriate control comparison for future experiments. RM expression upregulated Wnt and TGF-β signalling and downregulated genes related to ribosome biogenesis, and erythrocyte and megakaryocyte differentiation (**Figure 2G**).

### RM binds and induces m6A modifications on RNAs

Since RM and MKL1 have opposite effects on megakaryocyte maturation, we sought to determine whether RBM15 and its RNA binding function would be the major mechanistic contributor to the oncogenic conversion of MKL1 function. We therefore devised a multi-omic strategy to investigate RNA binding by RM and its effects on RNA modification, taking advantage of our FLAG-tagged models (**Figure 2A** and **2B**).

### RM binds common and unique targets with respect to RBM15

To compare transcripts bound by RM vs RBM15, we used anti-FLAG eCLIP-seq^23,24^ (Biancon, Joshi, et al., 2022; Van Nostrand et al., 2016) on HEL-RM and HEL-RBM15 cells with HEL-parental cells serving as negative controls. We identified 90,915 high-confidence binding sites of RM and/or RBM15 across 6,144 genes (**Figures 3A** and **S3A**), 60% of which (54,848 in 5,099 genes) were shared by RM and RBM15 (**Figure 3B**). Shared binding sites were enriched for pathways involving stress granules, negative regulation of erythrocyte differentiation, and RNA binding, processing, and transport (**Figure 3C**) and were enriched in coding and intronic sequences (**Figure S3B**). The high percentage of shared binding sites suggests that RM retains the RNA binding capacity of RBM15.

**Figure 3.**
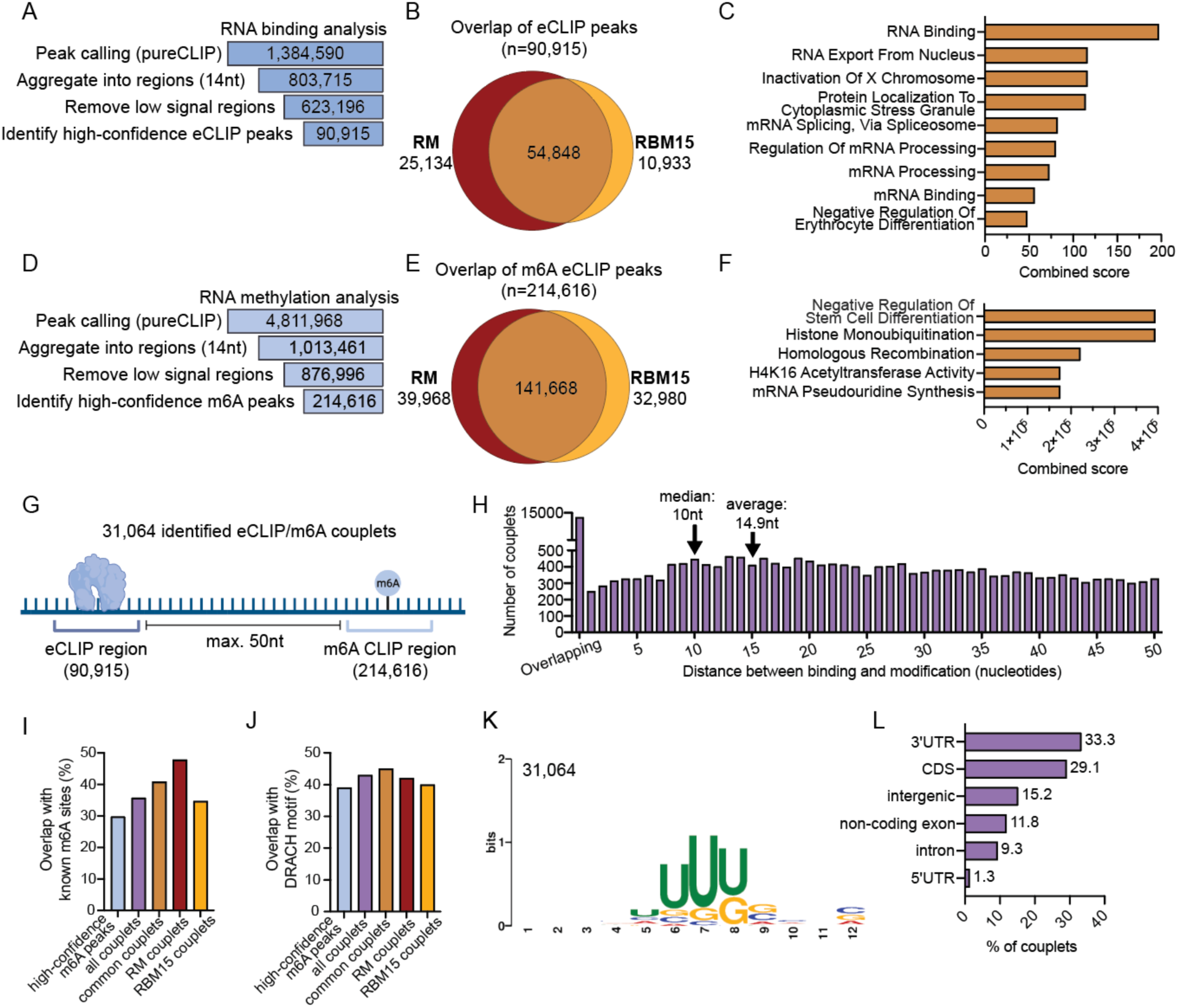
Integration of RNA binding and m6A modification identifies direct targets of RM and RBM15. (a) Steps in eCLIP experiment analysis pipeline; number indicates remaining sites after each filtering step. (b) Overlap of RM and RBM15 binding sites and (c) GO term analysis of shared sites. (d) Steps in m6A-eCLIP experiment analysis pipeline; number indicates remaining sites after each filtering step. (e) Overlap of m6A modified sites in HEL-RM and HEL-RBM15 cells and (f) functional enrichment analysis of shared m6A sites. (g) Schematic of couplets. (h) Distance between binding and modification in all couplets. Percentage sites that contain (i) a previously identified m6A site or (j) a DRACH motif in all regions, all high-confidence regions, common couplets, and RM- and RBM15-specific couplets. (k) Binding motif for all couplets. (l) Genomic location of all couplets. *See also Figure S3*.

Approximately 30% (25,134 sites in 1,496 genes) sites were differentially bound by RM, with associated genes enriched for pathways related to histone modification and Wnt and TNF signalling (**Figure S3C**). Genes differentially bound by RBM15 (10,933 sites across 1,195 genes) were enriched for cell cycle regulation and megakaryocytes/platelets (**Figure S3D**). Compared to RBM15, RM more frequently binds transcripts from intergenic regions (53% vs 20%) and less frequently from intronic regions (20% vs 44%) (**Figure S3B**). Genes differentially bound by RM and RBM15 tend to be upregulated by the respective proteins (RM: 35%, RBM15: 23%, **Figures S2E** and **S2F**), whereas of 5,099 commonly bound genes, only 9% are upregulated by RM and 6% by RBM15 (**Figure S3G**).

### RM partially alters the m6A RNA landscape with respect to RBM15

m6A eCLIP-seq on HEL-RM and HEL-RBM15 identified 214,616 high confidence m6A- modified sites in 19,253 genes (**Figures 3D** and **S3H**). m6A sites were enriched in the 3’UTR and CDS regions, and approximately 30% overlapped with previously identified^39,40^ m6A sites (**Figure S3I**). RM and RBM15 shared 66% (141,668 in 14,419 genes) of sites (**Figure 3E**) that were similarly localized to 3’UTR and CDS regions (**Figure S3J**) and were enriched for negative regulation of stem cell differentiation and histone modification (**Figure 3F**).

m6A sites identified only in HEL-RM (∼19%, 39,968 sites across 3,907 genes) were in genes enriched for suppression of apoptosis, m6A-containing RNA-binding, and AMKL (**Figure S3K**). m6A sites identified in only HEL-RBM15 (15%, 32,980 sites across 6,135 genes) were enriched in genes related to splicing and cell cycle regulation (**Figure S3L**). Compared to RBM15, RM-associated m6A modifications were more frequent at the 3’UTR (∼30% vs 19%) and less frequent in non-coding exons (∼9% vs 28%) (**Figure S3J**). Genes differentially modified were preferentially upregulated in respective cell lines (RM: 13.1%, RBM15: 4.0%) while few commonly modified genes had differential expression (**Figures S3M, S3N,** and **S3O**).

### Integration of RNA binding and m6A modification identifies direct targets of RM and RBM15

To determine which m6A sites are potentially modified due to RBM15 or RM binding, we integrated anti-FLAG eCLIP with m6A eCLIP to identify “paired” sites of binding and modification in close proximity.

### Paired binding and m6A modification “couplets” identify RM- versus RBM15-specific interactions with RNA

We considered m6A sites within 50nt ^41^ of RM and/or RBM15 binding regions to comprise a couplet. We identified 31,064 couplets (**Figure 3G**), which often contained overlapping binding and modification regions (average distance 14.9nt, **Figure 3H**). Couplets were enriched for previously published m6A sites^39,40^, for the DRACH motif and for a U-rich motif (similar to a published RBM15 binding motif^5^) and were primarily localized to 3’UTR and coding sequences (**Figures 3I, 3J, 3K**, and **3L**).

### RM differentially binds RNA and induces m6A modifications compared to RBM15

Of 31,064 couplets across 3,441 genes, 43% (13,372 couplets, 2,307 genes) were shared by RM and RBM15 and were enriched for regulation of transcription, chromatin remodeling, RNA processing, and RNA splicing pathways (**Figure S4A**), and 57% (17,692 couplets, 2,365 genes) were differentially targeted by RM or RBM15 (**Figure 4A**).

**Figure 4:**
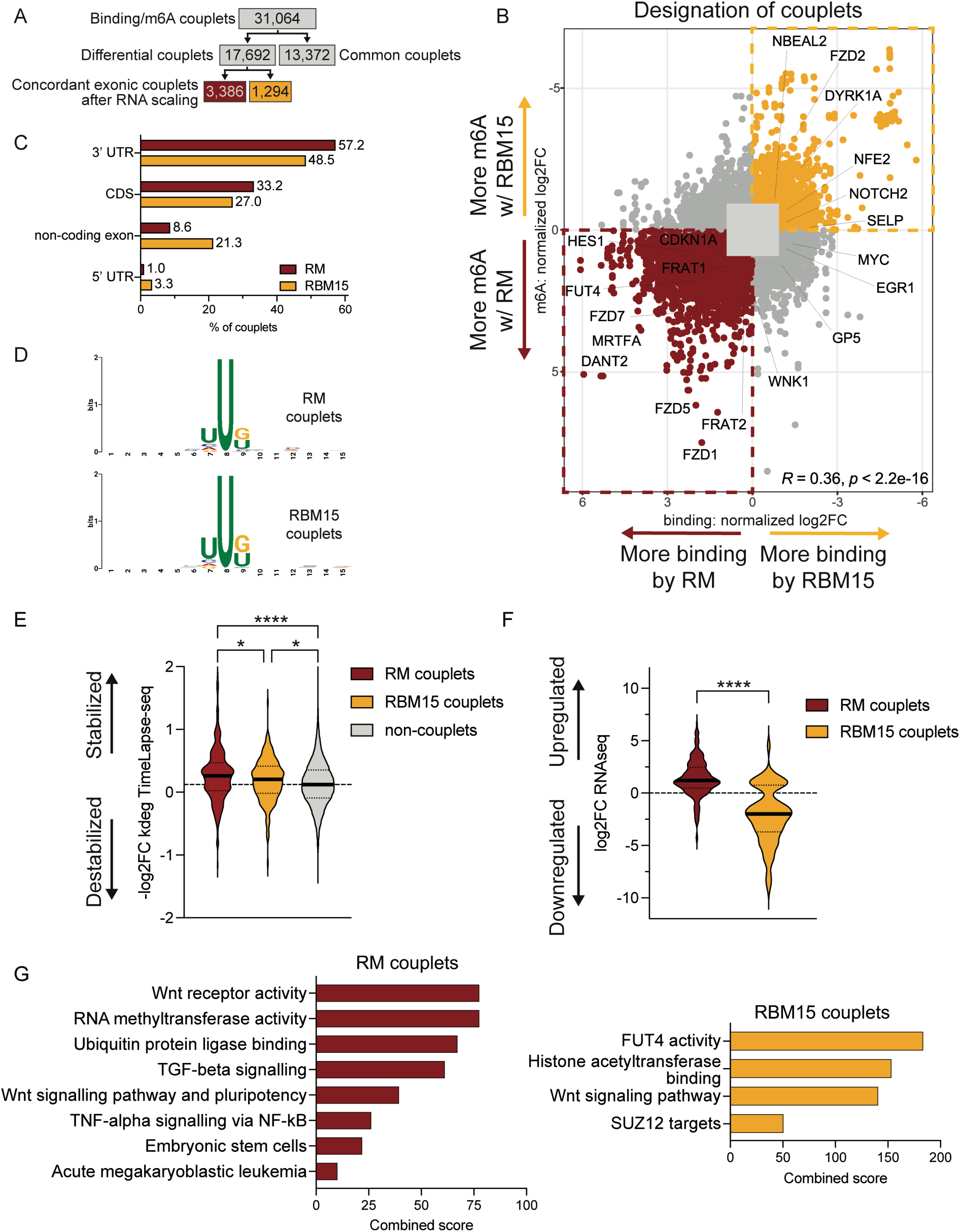
Integrated analysis of couplets, RNA-seq, and Timelapse-seq reveals RM-specific increases in expression and stability and identifies perturbations in the Wnt signalling pathway. **Multi-omics analysis reveals perturbations in the Wnt signalling pathway.** (a) Couplet identification pipeline; number indicates remaining sites after each filtering step. (b) Scatterplot identifying differential couplets after RNA scaling. Red box highlights RM-specific targets; yellow box highlights RBM15-specific targets. (c) Genomic location of differentially targeted couplets. (d) MEME Suite analysis identifies similar motifs in RM- and RBM15-specific couplets. (e) TimeLapse-seq shows RM couplets have increased stability compared to RBM15 couplets and non-couplets in RM-expressing cells (median indicated with solid line, quartiles indicated with dotted line; *p<0.05, ****p<0.0001, one-way ANOVA with Tukey’s multiple comparisons test). (f) Integration of RNAseq with RM- and RBM15-specific couplets shows differing expression outcomes (median indicated with solid line, quartiles indicated with dotted line, p<0.0001, independent t test). (g) Functional enrichment analysis of RM-specific couplets with increased expression in HEL-RM cells (top, red) and RBM15-specific couplets with decreased expression in HEL-RBM15 cells (bottom, yellow). *See also Figure S4*.

We observed an association between altered expression and altered binding and/or m6A modification (Pearson R=0.77, p=2.2e-16 and Pearson R=0.74, p<2.2e-16, respectively, **Figure S4B**). To increase confidence that differential binding was the primary event, we removed the potential influence of transcript abundance by normalizing couplet signals for RNA expression. For 10,634 out of 17,692 potential differential couplets, we could not exclude that differential binding/modification could be due to differential transcript abundance. We ruled out an effect of RNA abundance on the binding/modification for the remaining 7,058 couplets, which were therefore classifiable as the primary events (absolute L2FC>1 for binding or modification) (**Figures 4A** and **4B**). After correction for RNA abundance, there remained a statistically significant positive correlation between differential binding and modification for these 7,058 couplets (Pearson R=0.36, p<2.2e- 16, **Figure 4B**), suggesting that RBM15 and RM binding directly determine m6A modification. The majority of couplets were “concordant”, with differential binding and m6A modification simultaneously increased or decreased (**Figure 4B**). In concordant RM- and RBM15-specific couplets, binding and modification were within close proximity (average distance RM: 15.24nt, RBM15: 15.20nt) (**Figure S4C**). Concordant RBM15- and RM-associated couplets were localized to similar genomic regions, with the majority in the 3’UTR, and had similar U-rich binding motifs (**Figures 4C** and **4D**) suggesting that the fusion of MKL1 to RBM15 does not significantly alter the spatial constraints of or interfere with RNA binding and m6A deposition.

### Integrated analysis of couplets, TimeLapse-seq, and RNA-seq reveals RM-specific increases in RNA stability and expression and identifies perturbations in the Wnt signalling pathway

#### Genes differentially targeted by RM have increased stability

Because m6A modification regulates RNA fate^4–7^, we performed TimeLapse-seq^28^ (**Figures S4D, S4E, S4F,** and **S4G**) which identified significant changes in transcript stability (k_deg_) and synthesis (k_syn_) rates in HEL-RM and HEL-RBM15. Although RM and RBM15 both resulted in overall transcript stabilization relative to parental controls (**Figure S4H**), direct comparison between them revealed significant differential effects (**Figure S4I**). RBM15 increased the stability of RNAs related to megakaryocyte and erythrocyte signatures. In contrast, RM promoted stabilization of RNAs related to ribosome biogenesis as well as stem cell pathways.

Integration of changes in k_deg_ and k_syn_ reveals that when focusing on concordant couplets, RM-specific couplets mostly have increased k_syn_ in HEL-RM relative to HEL-RBM15 (**Figures S4J,** red dots, and **S4I**, black dots), whereas RBM15-specific couplets mostly have decreased k_syn_ in HEL-RBM15 relative to HEL-RM (**Figure S4K**, yellow dots), suggesting that transcriptional changes play a driving role in overall expression these genes. Because we observed an RM-specific effect on m6A deposition, we next focused on changes in stability. Consistent with context-dependent effects of m6A on RNA stability, both stabilization and destabilization of transcripts were promoted by RM and RBM15 (**Figures S4J** and **S4K**, red and yellow dots). In parental cells, genes associated with RM-specific couplets had shorter half-lives than non-couplet genes (**Figure S4L**). However, upon RM expression, RNAs associated with RM-specific couplets were significantly stabilized compared to RBM15 couplets and non-couplet genes (**Figure 4E**), suggesting there is m6A-mediated stabilization of RM target transcripts.

#### Genes differentially targeted by RM have increased expression and include oncogenic signatures

TimeLapse-seq showed both increased stability and synthesis of RM target genes, which corresponds to an overall increase in gene expression. While RM-specific couplets were more likely to be upregulated by RM, RBM15-specific couplets were more likely to be downregulated by RBM15 (**Figures 4F** and **S4M**). These findings suggest that differential targeting by either protein functions to change expression of the target genes and shed light on a potential mechanism of RM-AMKL. RM couplets that were upregulated by RM (176/488) were significantly enriched for potentially oncogenic signalling pathways including Wnt signalling (**Figure 4G**, red). RBM15 couplets that were downregulated by RBM15 (101/356) were also enriched for Wnt signalling as well as SUZ12 targets and FUT4 activity (**Figure 4G**, yellow). This suggests that fusion of RBM15 with MKL1 results in aberrant upregulation (and inability to downregulate) Wnt-related genes.

### METTL3 inhibition in murine RM-AMKL decreases Fzd expression

To determine whether the response of 6133 and CAOM cells to METTL3 inhibition was directly linked to the RM fusion protein, we queried genes that were differentially bound, m6A-modified, and upregulated by RM in HEL cells and were downregulated by METTL3 inhibition in 6133 and CAOM cells as candidate oncogenes in RM-AMKL. Approximately 71% of RM target genes (347/488) in HEL cells were also expressed in CAOM and 6133 cells; 27 were also downregulated by STM3675. These genes were greatly enriched for the Wnt pathway, including *Frizzled* genes, further implicating their role in RM-AMKL. The murine AMKLs express different m6A-dependent Fzd genes, suggesting convergent leukemogenic pathways; *Fzd8* was most highly expressed in 6133 cells, while *Fzd5* and *Fzd7* were more highly expressed in CAOM cells. STM3675 resulted in downregulation of these *Fzd* transcripts and their respective proteins (**Figures 5A, 5B, S5A,** and **S5B**). Additional *Frizzled* genes that were not directly bound and modified by RM were also upregulated in HEL-RM cells and downregulated in murine AMKLs by STM3675 (**Figure 5A**).

**Figure 5.**
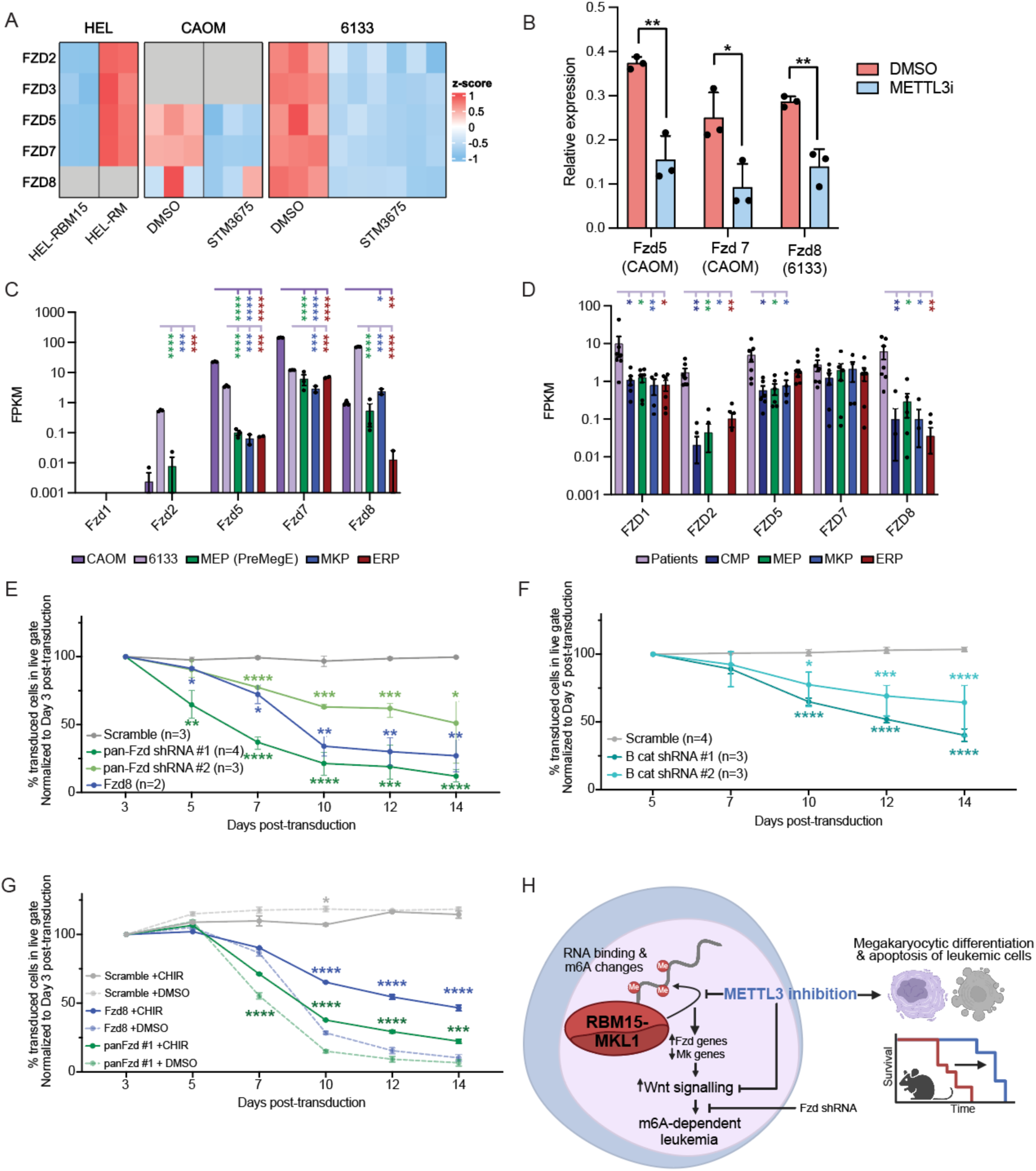
Wnt signalling is upregulated in RM-AMKLs and is required for cell growth. (a) Heatmap showing expression levels of Frizzled genes in all RNAseq experiments. (b) Protein levels of the most highly expressed Fzd genes in CAOM (Fzd5 and Fzd7) and 6133 cells (Fzd8) with Vehicle or METTL3 inhibitor treatment. (*p<0.05, **p<0.01, independent t tests). Compared to healthy control common myeloid progenitors (CMP, navy), megakaryocyte-erythroid progenitors (pre-MegE or MEP, green), megakaryocyte progenitors (MKP, blue) and erythroid progenitors (ERP, red), Frizzled genes are more highly expressed in (c) mouse models of AMKL (6133, purple and CAOM, dark purple) and (d) RM-AMKL patients (purple). Bars represent mean with SEM. 6133 cells transduced with lentivirus shRNA targeting (e) Fzd8 and panFzd (Fzd 1,2,5,7,8) and (f) β-catenin have decreased growth compared to scrambled control. (g) β-catenin agonist CHIR99021 partially rescues knockdown of Fzd1,2,5,7,8 and Fzd8 in 6133 cells. Error bars represent mean with range. (h) Model. For human samples, stars represent q-value after multiple Mann-Whitney tests with Benjamini-Hochberg correction for multiple comparisons. For mouse samples and knockdown experiments, stars represent q-value after multiple unpaired t-tests with Benjamini-Hochberg correction for multiple comparisons. (ns not indicated, *q<0.05, **q<0.01, ***q<0.001 , ****q< 0.0001). *See also Figure S5*.

### Wnt signalling is upregulated in RM-AMKLs and is required for cell growth

We determined the extent to which Fzd genes are differentially expressed in leukemias compared to primary nonmalignant progenitors by comparing relative mRNA expression of Fzd genes in 6133 and CAOM cells to murine primary cells from related stages of hematopoiesis^23^. In murine RM-AMKLs, Fzd genes are significantly more highly expressed than in pre-megakaryocyte erythroid progenitors (PreMegE), megakaryocyte progenitors (MkP), and erythroid progenitors (ErP) from healthy wildtype mice (**Figure 5C**). We repeated this comparison in human cells using published datasets^1,32^. Similar to the murine cell lines, pediatric RM-AMKL cells^1^ have higher levels of Frizzled RNA than related progenitor cells from healthy donors^32^ (**Figure 5D**). Similarly, a Wnt signalling score is increased in RM-AMKL patients compared to healthy progenitors (**Figure S5C**) suggesting that signalling downstream of Fzd proteins is important to RM-AMKL and that the mechanism by which they are upregulated in RM-AMKL may represent a potential therapeutic target. Interrogation of published datasets of pediatric AML^31^ revealed generally increased Wnt scores in other genotypes of AMKL compared to other AMLs and to healthy progenitors (**Figure S5C**), suggesting mechanisms converging on Wnt signalling may contribute to other genotypes of AMKL.

We tested whether targeting components of the Wnt signalling pathway, including *Fzd* and *β-catenin* (**Figure S5D** and **S5E**), affects AMKL growth. Using GFP-expressing shRNAs targeting single Fzd genes (*Fzd5, Fzd7*, and *Fzd8*) and pan-Fzd shRNAs (against complementary regions in *Fzd1, 2, 5, 7,* and *8*) revealed that single knockdown of *Fzd8* in 6133 cells caused a loss of transduced cells over time (**Figure 5E**, blue); pan-Fzd knockdown also caused cell loss (**Figure 5E**, green). Single knockdowns of *Fzd5* and *Fzd7* did not significantly affect 6133 cells (**Figures S5F** and **S5G**). shRNA mediated knockdown of *β-catenin* also resulted in loss of transduced cells over time (**Figure 5F**). Treatment of 6133 cells with a β-catenin agonist (GSK3β inhibitor CHIR-99021^42^), partially rescued the effect of pan-Fzd or Fzd8 knockdown (**Figure 5G**, solid lines), strongly suggesting that the decreased growth of 6133 cells after *Fzd* gene knockdown is due to decreased stabilization of β-catenin.

## Discussion

Despite well described functions of RBM15 and MKL1, their roles within the RM fusion protein remain undescribed. In this study, we determine the effects of RM on RNA processes and elucidate key transcriptomic and epitranscriptomic aspects of RM function. Our findings demonstrate that RM retains the RNA-binding and m6A modification abilities of wildtype RBM15 and additionally binds, modifies, stabilizes, and upregulates distinct RNAs not affected by RBM15.

While we show an increased stability of RM targets, there is also increased transcription of many RM targets, suggesting these genes are m6A-modified and stabilized in concert with an increase in synthesis. Investigation into the potential role of RM as a driver of gene expression will likely yield important information about the chromatin-associated effects of RM.

Because RBM15 is an integral part of the m6A writer complex, the RM fusion protein offers an opportunity to shed light on the role of m6A-mediated transformation in AMKL. Consistent with previous findings^20^, we show that targeting the m6A modification process may be an effective treatment for AMKL. METTL3 inhibition increased apoptosis and promoted differentiation of murine RM-AMKL cells and prolonged survival in transplanted mice. RNA-seq revealed downregulation of RM-specific targets, particularly Frizzled family genes, revealing a critical link between METTL3 and the WNT pathway, a well-established driver of various cancers^43^.

Our study reveals that RM directly interacts with Frizzled RNAs, leading to their upregulation. This is consistent with previous studies reporting upregulation of WNT ligands in CBFA2T3-GLIS2-driven AMKL^44–46^ and m6A-mediated stabilization/translation of Fzd RNAs contributing to oncogenesis^47–50^. In addition to the effect of m6A inhibition, reducing Frizzled expression via shRNA impairs murine AMKL cell growth, underscoring the critical role of downstream signalling in RM-AMKL pathogenesis.

Our newly described CAOM cells provide a robust *in vivo* model of murine RM-AMKL. While 6133 cells and CAOM cells share an MEP-like expression profile, CAOM cells also expressed genes such as *Spi1* and *Runx2* suggesting that CAOM cells were potentially immortalized at an earlier, more multi-potent cell state. This mimics patient heterogeneity and provides a new physiologically relevant model for AMKL research.

Several limitations should be acknowledged. First, we defined a “couplet” as binding and modification within 50nt^41^ which may exclude more distant modifications. Second, we focused on concordant exonic couplets; however, we also identified couplets that did not fit these criteria. For example, many non-exonic RM couplets are in DANT2, a long non-coding RNA thought to regulate constitutive heterochromatin at *DXZ4*^51^. Similarly, discordant couplets also merit further study. Third, TimeLapse-seq analysis may have been limited by low expression of some genes in HEL-RBM15 cells, restricting conclusions drawn from this dataset. Lastly, the lack of an ideal model for sequencing experiments necessitated use of HEL cells for multi-omic studies. The lack of a suitable control for murine RM-AMKL cells underscores the need for a new model for future studies.

In conclusion, our study delineates the transcriptomic and epitranscriptomic targets of the RM fusion protein elucidating a potential mechanism for RM-mediated leukemogenesis. This mechanism, whereby RM binding, m6A modification, and upregulation of Fzd genes drive dysregulated Wnt signalling in RM-AMKL (**Figure 5H**), represents an exciting new avenue for targeted therapy for RM-AMKL patients.

## Acknowledgements

We thank Martin Matthews and Yale’s Pathology Tissue Services for help with histology. We thank Evrett Thompson for assistance with RNA-seq analysis on primary human cells and Dr. Juliana Xavier-Ferrucio for assistance with RNA-seq analysis on primary murine cells. We thank Yale Flow Cytometry Core (supported in part by by an NCI Cancer Center Support Grant # NIH P30 CA016359) for their assistance. We thank the Yale Center for Genome Analysis (supported by the NIGMS of the NIH under Award Number 1S10OD030363-01A1) for their assistance with sequencing.

M.Y.M. has received funding from the Lo Graduate Fellowship for Excellence in Stem Cell Research and the NIH via NCI F31CA27157 and a pilot grant from the Cooperative Center of Excellence in Hematology (CCEH) (NIDDK U54DK106857). G.B. has received funding from the ASH Scholar Award and the Edward P. Evans Foundation. M.W. has received funding from the Leukemia and Lymphoma Society. C.R. has received funding from the NRRP (National Recovery and Resilience Plan) CUP E63C22001220001. M.D.S. has received funding for TimeLapse-related studies from the NIH via NIGMS award number R01GM137117. T.T. has received funding from the AIRC Foundation for Cancer Research under MFAG 2020 (ID. 24883 project). S.H. has received funding from the NIH via NCI award numbers R01CA222518, R01CA253981, and 1R01CA266604. D.S.K. has received funding from the NIH via NCI R01CA222518 and NIDDK RC2DK122376.

## Author Contributions

Conceptualization – MYM, GB, SH, DSK

Data curation – MYM, GB, CR, IM, AP-U, TT

Formal Analysis - MYM, GB, MW, CR, IM, AP-U, MDS, TT

Funding acquisition – MYM, MW, TT, SH, DSK

Investigation - MYM, GB, MW, AP-U, LW, JAE, YO-R, OR, SH

Methodology – MYM, GB, MW, CR, IM, AP-U, MDS, TT, SH, DSK

Project administration – MYM, GB, MW, TT, SH, DSK

Resources – TT, SH, DSK

Software – GB, CR, AP-U, MDS, TT

Supervision – GB, MDS, TT, SH, DSK

Validation – MYM, GB, MW, CR, IM, AP-U, LW, JAE, TT

Visualization – MYM, GB, MW, CR, IM, AP-U, YO-R, OR, TT, SH, DSK

Writing – original draft - MYM, GB, MW, CR, AP-U, TT, SH, DSK

Writing – review & editing – MYM, GB, MW, CR, IM, AP-U, LW, JAE, MDS, YO-R, OR, TT, SH, DSK

