## Supplemental Methods for "RBM15-MKL1 fusion protein promotes leukemia via m6A methylation and WNT pathway activation"

#### RESOURCE AVAILABLITY

**Lead contact**

**Materials availability**

Plasmids and cell lines generated in this study are available from Diane S. Krause and Stephanie Halene with completed Materials Transfer Agreements. Additional details related to the chemical characterization of STM3675 can be found in the relevant published patent (International Publication Number: WO2021/111124 Al) at World Intellectual Property Organization.

**Data and code availability**

Sequencing data generated in this study are publicly available and accessible as of the date of publication on Gene Expression Omnibus (GEO) under GEO: GSE######
Any additional information required to reanalyze the data reported in this paper is available from the lead contact upon request.

#### EXPERIMENTAL MODEL AND SUBJECT DETAILS

**Cell Lines**

Human erythroid leukemia (HEL) cells were stably transfected with pcDNA6/TR and were selected for with 5µg/mL blasticidin. These cells represent the parental control cell line. pcDNA6/TR cells were then additionally transfected with pcDNA5/TO-FLAG-RBM15 or with pcDNA5/TO-FLAG-RBM15-MKL1. Cells were maintained in RPMI with 10% tetracycline-free fetal bovine serum (FBS), 1% penicillin/streptomycin, and 1% L-glutamine. PCDNA5 cells were additionally maintained with 375µg/mL hygromycin. For induction of transgenes, cells were treated with 10ng/mL doxycycline for 24 hours. To induce differentiation, $2x10^5$ cells per mL were cultured in the presence of 15nM 12-O-tetradecanoyl-phorbol 13-acetate (TPA; Sigma-Aldrich, St Louise, MO) for 24 or 72 hours.

6133 cells were a kind gift of Thomas Mercher and Gary Gilliland and were maintained in RPMI with 10% FBS, 1% penicillin/streptomycin, 1% L-glutamine, 100ng/mL murine stem cell factor (mSCF, ConnStem S2000), and 10ng/mL murine interleukin 3 (mIL3, ConnStem I2003) [1]. The CAOM megakaryoblastic leukemia cell line was generated from a Rbm15-MKL1 knockin mouse that spontaneously developed leukemia as previously described.[1] CAOM cells carry flox sites in the *Cebpa* locus from a previous experimental crossing to C57BL/6-*Cebpa*$^{tm1Dgt}$/J but do not express Cre recombinase and are propagated via serial transplantation in Pep3B (B6.129S6(CBA)-*Cebpa*$^{tm1Dgt}$/J) recipients. CAOM cells are grown in DMEM, 10% FBS, 1% penicillin/streptomycin, 1% L-glutamine, 100ng/mL mSCF and 10ng/ml mIL-3 (ConnStem Cat# I2003).

**In vivo leukemia model**

All mice were bred and maintained under specific-pathogen-free conditions at the animal facility of Yale University School of Medicine. Animal experiments were performed under protocols approved by the Institutional Animal Care and Use Committee of Yale University. Both female and male mice were used in experiments.

Syngeneic B6.SJL-*Ptprc*[a] *Pepc*[b]/BoyJ (Peb3b, JAX: 002014) transplant recipient mice were 6-8 weeks of age. CAOM cells were derived from progeny of crossing of Cebpa$^{f/f}$ (B6.129S6(CBA)-Cebpatm1Dgt/J; JAX: 006447) to *Rbm15-MKL1* knockin mice without deletion of the *Cebpa* gene. 400,000 CAOM cells from spleens of mice carrying CAOM leukemia were transplanted into sub-lethally irradiated (600 cGy) Pep3b mice. To ensure the reproducibility of data, we stratified animals to Vehicle versus STM3675 treatment by integrating the percentage of c-kit+ cells and platelet counts in peripheral blood of transplanted mice 14 days after transplantation of CAOM cells and started treatments on D17 after transplantation. After stratification to treatment versus vehicle mice were gavaged daily 6 out of 7 days per week with 100mg/kg STM3675 for a total of 17 days. Animals were monitored twice daily and euthanized when moribund and analyzed for engraftment in bone marrow and spleen via histology. Animals who succumbed to leukemia were not further analyzed. Spleens were weighed at time of euthanasia as surrogate for leukemia engraftment levels.

### METHOD DETAILS

#### METTL3 inhibitor – characterization of STM3675

*METTL3/14 RapidFire Mass Spectrometry Methyltransferase Assay*

RapidFire assay was performed as previously described[2].

*SPR Assay*

SPR assay was performed as previously described[3].

*Electroluminescence-Based ELISA for the Quantification of m6A Modification on PolyA+ RNAs*

Assay was performed as previously described[3].

*Pharmacokinetic (PK) profiling and in-vivo PK-PD correlation*

For PK profiling, 3 male C57Bl/6J mice were orally dosed with a single dose of STM3675 dissolved in a buffer containing Tween80 / Captisol 10% in Acetate buffer pH4,6 50mM (5%/95%) to generate 10 mg/mL solution. A volume of 10 mL/kg was used. At the indicated times, tail vein microsampling was performed and the concentration of STM3675 was determined using standard Mass Spectrometry methods.

For PK-PD correlation, naïve mice were administered a single dose of STM3675 at the indicted dose levels. 7 hours after treatment, blood and spleen samples were then taken for PK and PD analyses, respectively. STM3675 level was determined using standard Mass Spectrometry methods. m6A quantification on polyA-RNA isolated from spleens was performed using the ECL ELISA method described in Guirguis et al[3].

#### METTL3 inhibitor – in vivo and in vitro experiments

For *in vitro* treatment of 6133 and CAOM cells, STM3006 and STM3675 stock solutions were generated at 10mM in DMSO diluted further to achieve final concentrations as indicated. For *in vivo* use STM3675 stock solution was prepared at 10mg/ml in Captisol/Citrate buffer pH3.0 and administered at 100mg/kg once daily via oral gavage for the indicated time points.

**Cell toxicity assay and morphologic analysis**

Cells were plated in 96-well plates in triplicate at 5,000 cells per well (6133) or 100,000 cells per well (CAOM) and treated with vehicle or serial concentrations of STM3006 and STM3675, as indicated, in 100µL DMEM/10% FBS/ Pen-Strep supplemented with recombinant murine Scf (100ng/ml) and IL-3 (10ng/ml). Media and drugs were refreshed on day 3 and on day 5, plates were measured using CellTiter-Glo® (Promega) per manufacturer's instructions and relative cell numbers normalized to vehicle. The IC50 was calculated by nonlinear regression using GraphPad Prism version 9.0. For morphologic analysis, cells were treated for 24, 48 and 72 hours at 1µM and ~200,000 cells were dropped onto Superfrost Plus™ Slides and allowed to settle for 15 minutes. Slides were washed 1x with PBS and fixed in methanol for 10 minutes, followed by Wright Giemsa staining per manufacturer's protocol (Siemens Healthineers, Cat# 06689653). Images were acquired on a Keyence BZ-X800 microscope (KEYENCE Corporation of America, Itasca, IL, USA). Cell size was analyzed and quantified using ImageJ [4].

**Flow cytometric analysis**

Ploidy of HEL and HEL-RM cells was determined using Propidium Iodide as described[5]. To study the block in HEL-RM cell maturation, the following antibodies were used to stain cell surface markers: CD49b-PE (BioLegend) and CD235A-APC (BioLegend). Flow cytometric analysis was performed with an LSRII flow cytometer (BD Biosciences) and FlowJo software (TreeStar, Ashland, OR).

Flow cytometry analysis was performed as described previously[6]. To characterize 6133 and CAOM cells, cells were blocked with rat anti-mouse CD16/32 antibody for 10 min, followed by staining with indicated antibodies in FACS buffer (2% BSA, 2 mM EDTA in PBS) in the dark at 4 °C for 20 min. To study apoptosis in 6133 and CAOM cells, 0.5 $x10^6$ cells were stained with the Annexin V Apoptosis Detection Kit following the supplier provided protocol (Biolegend, Cat#420403). To determine differentiation of 6133 and CAOM cells, the following antibodies were employed: 7-AAD (Biolegend), ckit - APC-Cy7 (Biolegend) combined with either CD41-FITC (Biolegend) or CD61-FITC (Biolegend). Flow cytometric analysis was performed on a FACSymphony (BD Biosciences) instrument. Flow data were analyzed with FlowJo software version v10.10 (TreeStar, Ashland, OR, USA)

**Histologic analysis**

Tissues were fixed in 10% Neutral Buffered Formalin (Fisher HealthCare, Cat #316-155). Femurs were further decalcified with Formic Acid Bone Decalcifier (Decal Chemical, NY, USA), washed in 70% ethanol, and tissues were embedded in paraffin and tissue sectioned at 5 µm and stained with hematoxylin & eosin. Images were acquired on a Keyence BZ-X800 microscope.

**RNA-seq**

For HEL cell RNA sequencing, RNA was isolated using the RNeasy Mini Kit (QIAGEN, cat# 74104) per vendor supplied protocol. For 6133 and CAOM RNA sequencing, cell pellets were dissolved in TRIZOL (Life Technologies, Cat #15596018) and prepared according to vendor supplied protocol. RNA underwent poly-A selection or rRNA depletion and reverse transcription, library preparation using the KAPA mRNA HyperPrep kit, and 100bp paired-end sequencing on the Illumina NovaSeq 6000 at the Yale Center for Genome Analysis.

For RNA-seq of murine progenitors, murine bone marrow was sorted as previously described[7,8]. Briefly, megakaryocyte progenitors were identified as $Lin^{-}cKit^{+}Sca^{-}CD150^{+}CD41^{+}$ and erythroid progenitors as $Lin^{-}cKit^{+}Sca^{-}CD150^{+}FcgRI/II^{-}CD105^{+}$. Bulk RNA-seq was performed on sorted progenitor populations. Libraries were prepared and sequencing conducted by the Yale Stem Cell Center Genomics Core facility on the Illumina HiSeq 4000.

**eCLIP-sequencing**

eCLIP-seq experiments were performed in triplicate according to previously published protocols[9–11] with the following adaptations. After 24h dox, HEL cells expressing FLAG-tagged RBM15 or RM or empty vector control were UV-crosslinked twice at 2000mJ/cm$^2$ using a Stratalinker at 254nm. Cells were sonicated for a total of 60 seconds in eCLIP lysis buffer and treated with RNAse 1 (ThermoFisher SCIENTIFIC cat# AM2295) for 3 minutes at 37ºC. Protein-RNA complexes were immunoprecipitated using Dynabeads (ThermoFisher SCIENTIFIC cat# 10004D) and 12ug anti-FLAG antibody (Sigma Aldrich, cat# F1804). Libraries were prepared as described[10,11]. Resultant libraries underwent 100bp paired-end sequencing on the Illumina NovaSeq 6000 at the Yale Center for Genome Analysis.

**m6A eCLIP-sequencing**

After 24h dox, 10 x 10^6 cells HEL cells expressing FLAG-tagged RBM15 or RM were pelleted, flash-frozen, and stored at -80º. RNA was extracted using QIAGEN RNeasy Mini kit with DNAse treatment (QIAGEN, cat# 74104) and quantified using nanodrop. mRNA was isolated from 75ug total RNA by oligo-dT beads (ThermoFisher Dynabeads™ mRNA Purification Kit, cat# 61006) and eluted in 5µL DNase-RNase free water. This was repeated until >8ug mRNA was obtained per sample. RNA was fragmented using 2 µL of RNA fragmentation reagent (ThermoFisher, cat# AM8740) at 75ºC for 6 minutes and RNA integrity and fragment size was assessed using an Agilent TapeStation. mRNA was incubated with 2µL of ribonuclease inhibitor (SuperaseIn) and 5ug anti-m6A antibody (202 011, Synaptic Systems) for 2 hours, rotating, at 4ºC. Each sample was transferred to a single well of 12-well plate on ice and crosslinked twice at 150mJ/cm$^2$ in a Stratalinker at 254nm. Antibody-RNA complexes were immunoprecipitated using Dynabeads (ThermoFisher SCIENTIFIC cat# 10004D). We then performed eCLIP as described above. Resultant libraries underwent 100bp paired-end sequencing on the Illumina NovaSeq 6000 at the Yale Center for Genome Analysis.

**TimeLapse-seq**

TimeLapse-seq experiments were performed in triplicate essentially as described previously[12]. HEL cells expressing FLAG-tagged RBM15, RM, or empty vector control were treated with dox for 24h. During the last 2h of dox treatment, HEL cells were supplemented with 100µM 4-thiouridine. Single replicate controls not labeled with 4-thiouridine were also collected for each condition. After 2h of 4-thiouridine treatment, HEL cells were fully resuspended and transferred into ice-cold PBS. Cells were pelleted for 5 mins at 500Xg and subsequently resuspended in 1mL of TRIzol reagent. Total RNA was isolated and extracted with phenol-chloroform. Following precipitation in isopropanol supplemented with 1mM DTT and 20μg of glycogen, RNA pellets were washed twice with fresh 75% EtOH. Dried RNA pellets were resuspended in DEPC-treated water and treated with Turbo DNase to deplete genomic DNA. After purification with one equivalent volume of Agencourt RNAClean XP beads, 5μg of eluted RNA were processed for TimeLapse chemistry using meta-chloroperoxybenzoic acid as the oxidant[12]. For each sample, 10ng of TimeLapse-treated RNA input were used to prepare sequencing libraries from the Clontech SMARTer Stranded Total RNA-Seq kit (Pico Input) with ribosomal cDNA depletion. Paired-end 100bp sequencing was performed on the Illumina NovaSeq 6000 at the Yale Center for Genomic Analysis.

**Immunoblot**
Cell pellets were lysed in RIPA buffer with proteinase inhibitor (Roche cat# 11836153001) and protein concentration was quantified using a Bradford assay (Bio-Rad cat# 5000006). Samples were boiled to denature proteins and separated in Mini-PROTEAN® TGX Stain-Free™ Protein Gels (Bio-Rad). Lysates were transferred to 0.45 μm nitrocellulose membranes with a standard wet transfer system at 30V for 90 minutes or at 100V for 90 minutes. Membranes were blocked with 5% skim milk in TBST for 60 min and incubated with primary antibodies overnight at 4˚C. Excess antibody was washed away with TBST (50 mM Tris pH 8.0, 150 mM NaCl, 0.1% Tween 20) 3 times. Membranes were incubated with HRP-linked secondary antibody diluted in 5% skim milk for 1hr at room temperature. After 3 washes, membranes were developed with SuperSignal™ West Femto Maximum Sensitivity Substrate (ThermoFisher Scientific, Cat# 34095).

**Plasmids and viral transduction**
Lentiviral constructs for shRNAs were ordered from VectorBuilder using the following target sequences:

Fzd8: GACCAGGCAGATGCCTTAAAT
panFzd #1: GGCCAGCTCCATCTGGTGGGT
panFzd #2: CATGCTCAAGTACTTCATGTG and CGCATCCGCACCATCATGAAG
β-catenin #1: GCGTTATCAAACCCTAGCCTT
β-catenin #2: TCTAACCTCACTTGCAATAAT
Fzd5: CGAGGTTCTGTGTATGGATTA
Fzd7: TGGAGCCCAGATGGGTTAATT
Scramble: CCTAAGGTTAAGTCGCCCTCG

All constructs were confirmed by Sanger sequencing.
For virus production, HEK293T cells were maintained in DMEM with sodium pyruvate, 10% FBS, 1% penicillin/streptomycin, 1% L-glutamine. For lentivirus production, media was supplemented with 1.1g/mL BSA, cells were transfected with constructs and FuGENE (FuGene cat # F6-1000), virus-containing media was harvested. For transduction, cells were incubated with virus and 8 μg/mL polybrene and were assessed for transduction efficiency on day 3 post-transduction.

**QUANTIFICATION AND STATISTICAL ANALYSIS**
**Bulk RNA-Seq analysis**

After quality control (FastQC version 0.11.9, https://www.bioinformatics.babraham.ac.uk/projects/fastqc/ ), reads generated from each sample were deduplicated with FastUniq (version 1.1, FastUniq download | SourceForge.net) and aligned to the mouse genome (GRCm39) or the human genome (GRCh38) with STAR (version 2.7.10a, - quantMode GeneCounts), using the Gencode M31 (mouse) and 37 (human) gene annotation, respectively. Normalization with the TMM method and identification of differentially expressed genes were performed with the edgeR (version 4.2.0) package in Bioconductor (https://bioconductor.org/). Differentially expressed genes were identified with the "glmQLFTest" function, using a double threshold on gene expression changes and associated statistical significance (absolute log2 fold change > 0.75, FDR < 0.05).

**eCLIP and m6A eCLIP-seq analysis**

Initial peak calling was performed with PureCLIP[13] (version 1.3.1, https://github.com/skrakau/PureCLIP). Nucleotides with a Hidden state of 3 (enriched + crosslink) were considered potential crosslink nucleotides. The regions of the initial crosslinked nucleotides were extended by 7 nucleotides upstream and downstream, resulting in 15 nucleotide long regions. Overlapping regions on the same strand were merged with the merge Bedtools function. Read coverage on each region for each replicate was quantified with Bedtools coverage. Only regions with a minimum count of 10 among all the replicates in at least 1 condition were considered as regions with sufficient signal. Additionally, for m6A eCLIP, only regions containing at least one A were considered.

Read counts for each region were normalized and differential analysis was performed in the same manner as RNA-seq. To identify high-confidence CLIP regions for eCLIP, only regions enriched over both input and control conditions were considered; for m6A eCLIP, only the regions enriched over input were considered (p-value <0.05 and log2 fold enrichment >1).

Differential analysis was performed to find differential meCLIP and eCLIP RM regions using meCLIP and eCLIP RBM15 as controls. Differentially bound or methylated regions were identified with EdgeR (p-value < 0.05).

To identify "couplets", we paired each binding site with the nearest m6A site (distance threshold: 50 nucleotides). Differential couplets were identified as couplets with p-value <0.05 for either eCLIP or m6A eCLIP (RM vs RBM15 comparison). Delta delta couplets were identified among differential couplets by first scaling RM vs RBM15 fold changes on RNA fold changes, and then selecting couplets with a resulting scaled fold change ≥ 1 or ≤ -1 in either eCLIP or m6A eCLIP.

To identify regions containing known m6A sites, regions were overlapped with known m6A sites retrieved from m6A-Atlas[14] (version 2.0, http://rnamd.org/m6a/index.php) and RMBase[15] (version 3.0, https://rna.sysu.edu.cn/rmbase3/).

Functional enrichment analysis was performed with Enrichr[16] (version 3.2, https://maayanlab.cloud/Enrichr/).

**TimeLapse-seq analysis: quantification and analysis of differential gene expression kinetics**

TimeLapse-seq reads were trimmed and aligned with HISAT-3N to the human hg38 genome as described in the TimeLapse preprocessing pipeline [12]. After T-to-C mutation calling, bakR and DESeq2 were used to calculate differences in kinetic parameters and gene expression levels across RBM15 overexpression, RM overexpression, and parental control conditions[17,18]. Library size factors were estimated using DESeq2. The cB output file of the TimeLapse preprocessing pipeline was used as a count table in bakR to model transcript-specific changes in RNA degradation and synthesis rates. bakR was run using the hybrid model with all read filtering parameters set to default. Transcripts with $P_{adj} < 0.05$ in $k_{deg}$ across conditions were called as significantly differentially stable. Analogously, transcripts with $P_{adj} < 0.05$ in $k_{syn}$ across conditions were called as significantly differentially synthesized. Mature RNA half-lives were calculated using the following equation: $t_{½} = \ln(2)/k_{deg}$.

**GO enrichment analysis**

Destabilized genes determined by TimeLapse-seq were further analyzed for GO enrichment analysis on WebGestalt (http://www.webgestalt.org), using Over-Representation Analysis (ORA) as method to define the biological process and enriched KEGG pathways.

**RNA-Seq data analysis of AMKL and AML patient samples**

RNA-Seq gene counts from AMKL, AML and healthy control samples were obtained from publicly available datasets[19–21]. Normalization of raw gene counts was performed with the TMM method in the edgeR (version 4.2.0) package in Bioconductor (https://bioconductor.org/). FPKM values were obtained with edgeR using Gencode 37 (human) gene annotation. UMAP dimensional reduction was performed based on FPKM values of genes within the Wnt signalling pathway, as annotated in BioPlanet (https://tripod.nih.gov/bioplanet/detail.jsp?pid=bioplanet_230&target=pathway).

Wnt pathway activation scores for each sample were obtained with the accordion_custom function from the cellmarkeraccordion package (version 0.9.0) (https://github.com/TebaldiLab/cellmarkeraccordion)[22].

**Statistical Analysis**

All statistical analyses were performed using GraphPad Prism version 9 (GraphPadSoftware) or R.

### Supplemental figure titles and legends

Figure S1

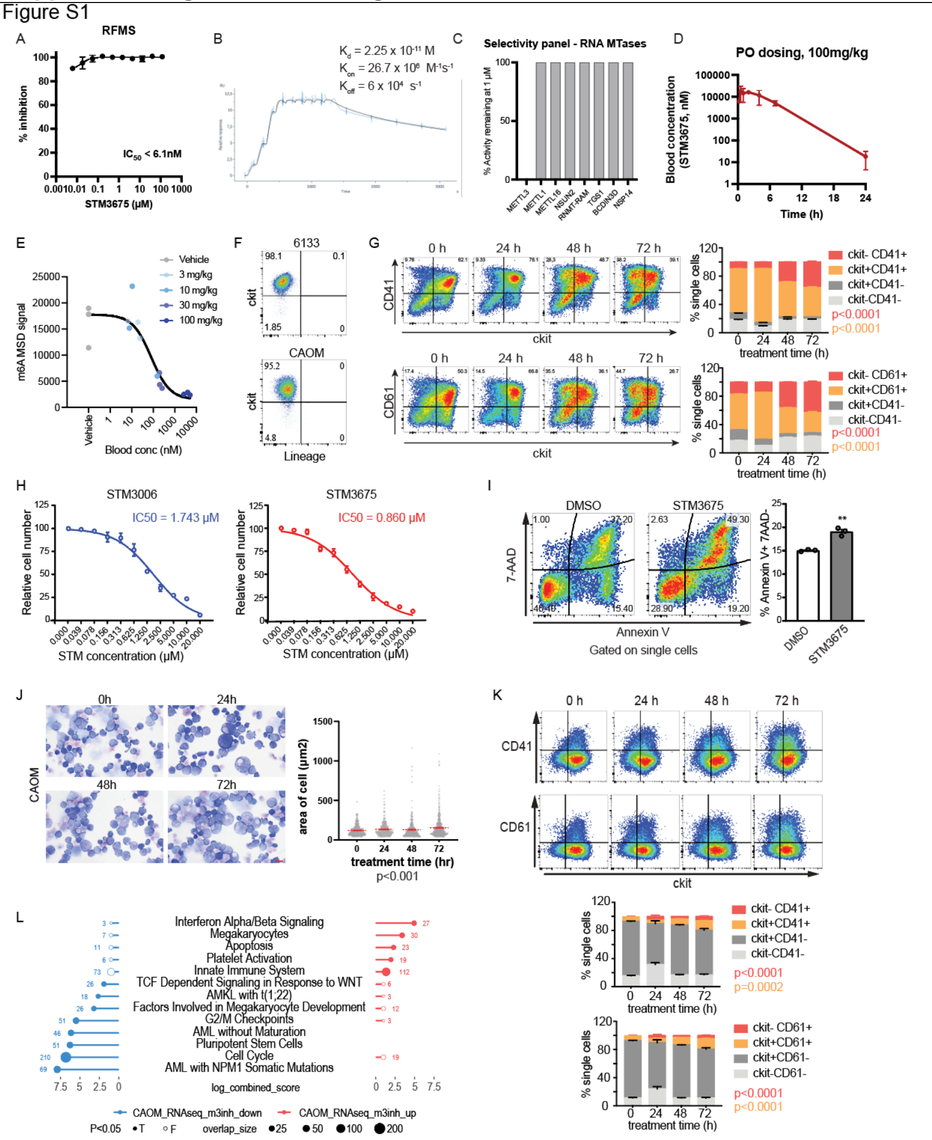


**Figure S1. Inhibition of Mettl3 induces AMKL blast death and prolongs survival in leukemic mice, Related to Figure 1.** (a) Inhibition of METTL3/14 enzymatic activity by STM3675 as determined by RapidFire mass spectrometry (RFMS), with IC50 depicted. (b) Binding affinity of STM3675 with METTL3/14 as determined by SPR in single-cycle kinetic mode. RU, relative units. (c) Remaining enzymatic activity following STM3675 treatment in a panel of 8 RNA methyl transferases, as measured by RFMS. (d) STM3675 *in vivo* pharmacokinetic profile. Total blood concentration was measured after a single PO dosing at 100 mg/kg in n=3 mice. (e) PK-PD correlation in mice treated with STM3675 at the indicated dose levels. Samples were taken 7 hrs after dosing. (f) Flow-cytometric characterization of 6133 cells assessing expression of CD41 and non-megakaryocytic lineage markers including CD5, CD11b, CD45R, Ly-6B.2, Gr-1 and Ter-119. (g) Flow-cytometric assessment and quantification of c-kit, CD41 and CD61 expression in 6133 cells treated with vehicle (0 hour) or STM3675 (1μM) for 24, 48 or 72 hours. A representative experiment of three is shown, p-value calculated by one-way ANOVA with Welch's test. (h) STM3006 and STM3675 are toxic to CAOM cells *in vitro* with IC50s of 1.74μM and 0.860μM, respectively. Total cell number was normalized to vehicle. IC50 was calculated by fitting toxicity curve using a nonlinear method. n = 3 biological replicates. (i) Flow-cytometric assessment and quantification of apoptosis (Annexin V + 7AAD -) in response to vehicle or STM3675 (1μM). A representative experiment of three is shown. Data are represented as mean ± SEM; the p-values were calculated using independent two-tailed Student's t test. **$p < 0.01$. (j) Wright-Giemsa staining and quantification of cell area ($\mu m^2$) of CAOM cells treated with Vehicle (0 hour) or STM3675 for 24, 48, 72 hours. Scale bar, 10 μm. p-values calculated with one-way ANOVA with Welch's test, $p < 0.001$. (k) Flow-cytometric assessment and quantification of c-kit, CD41 and CD61 expression in CAOM cells treated with vehicle (0 hour) or STM3675 (1μM) for 24, 48 or 72 hours. A representative experiment of three is shown, p-value calculated by one-way ANOVA with Welch's test. (l) Functional enrichment analysis of differentially expressed genes from CAOM cells treated with STM3675 or vehicle (1,846 upregulated genes and 1,692 downregulated genes).

Figure S2

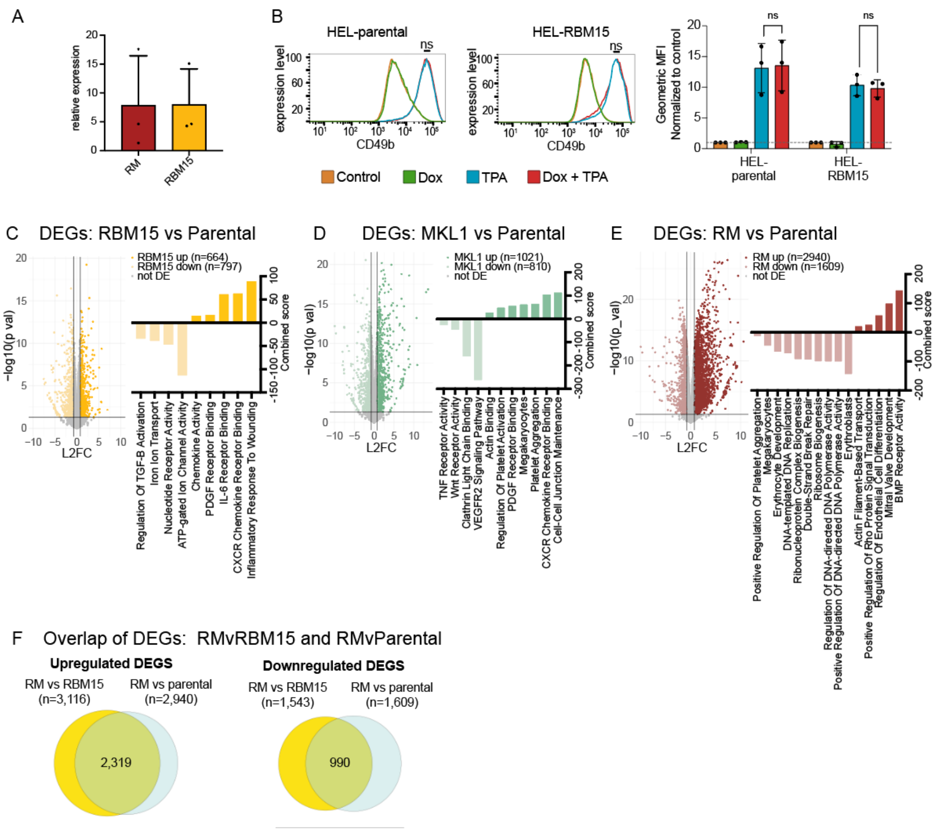


**Figure S2. Additional information about the HEL cell model, Related to Figure 2.** (a) Quantification of n=3 western blots of RM (red) or RBM15 (yellow) expression relative to parental controls. (b) Flow cytometry on HEL-parental (left) and HEL-RBM15 (right) without dox (orange), with dox (green), with TPA (blue), and with dox and TPA (red) shows no change in expression of megakaryocyte maturation marker CD49b in HEL-RBM15 cells treated with dox and TPA (red) compared to TPA alone (blue), data represented as percentage of max. Differential expression analysis of RNAseq from (c) HEL-RBM15 vs parental and (d) HEL-MKL1 vs parental and (e) HEL-RM vs parental. (e) overlap of DEGs from RM vs RBM15 and RM vs parental (right panel = upregulated DEGs, left panel = downregulated DEGs).

Error bars represent mean ± SD, n=3, relevant p values are plotted (ns not significant, 2-way ANOVA with Fisher’s LSD).

Figure S3

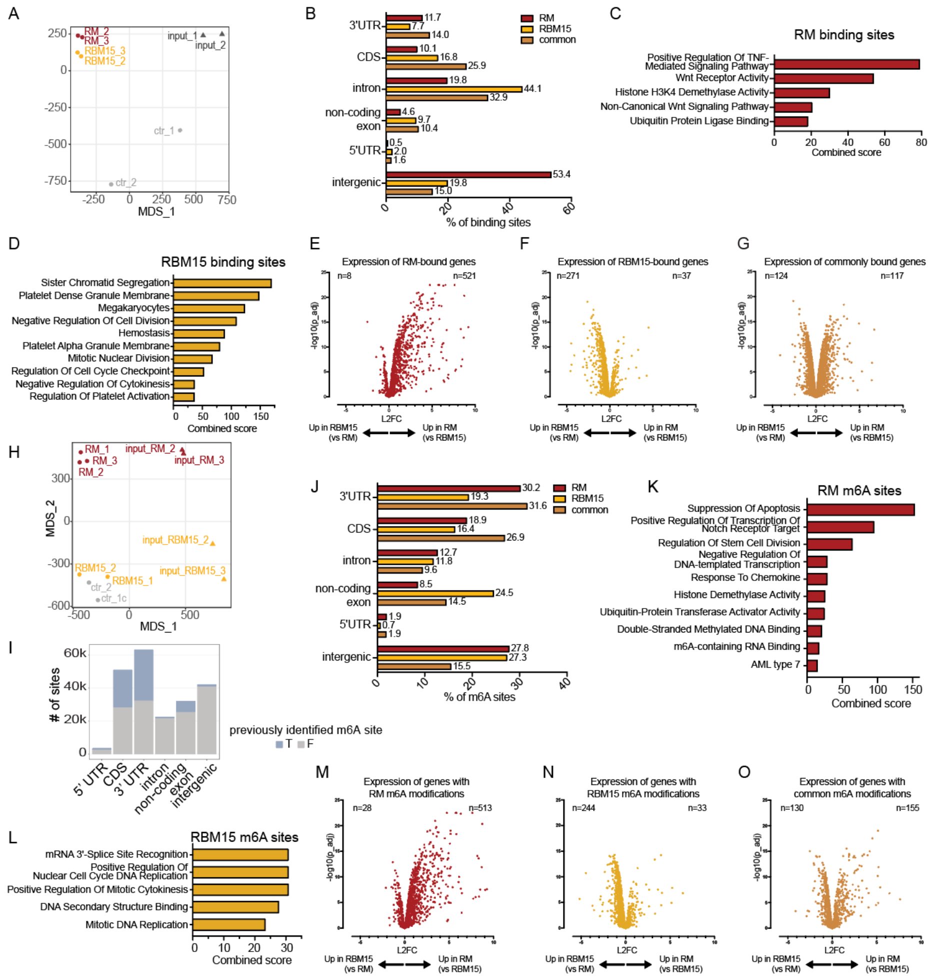


**Figure S3. Supplemental information about eCLIP and m6A eCLIP experiments, Related to Figure 3.** (a) MDS plot of anti-FLAG eCLIP-sequencing experiments showing clustering of replicates. (b) Location of RM (red), RBM15 (yellow), and common (orange) binding sites. Functional enrichment analysis of (c) RM-specific and (d) RBM15-specific binding sites. RNA-seq expression levels of (e) RM-specific, (f) RBM15-specific, and (g) commonly bound genes. (h) MDS plot of anti-m6A eCLIP-sequencing experiments showing clustering of replicates. (i) Frequency of previously identified m6A sites with breakdown of genomic location in HEL cell anti-m6A eCLIP experiments (previously identified = blue). (j) Location of RM (red), RBM15 (yellow), and common (orange) m6A sites. Functional enrichment analysis of (k) RM-specific and (l) RBM15-specific m6A sites. RNA-seq expression levels of (m) RM-specific, (n) RBM15-specific, and (o) common m6A sites.

Figure S4

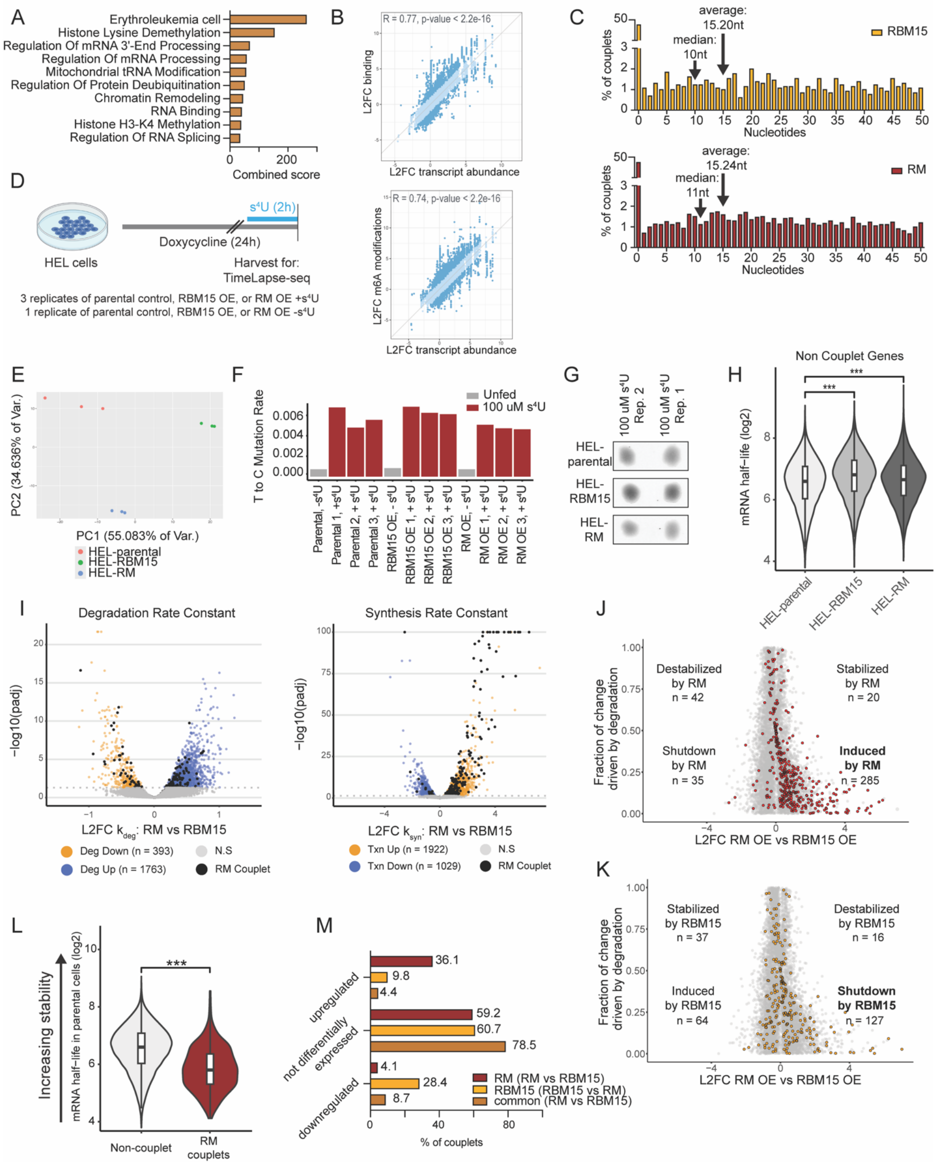


**Figure S4. Additional analyses of couplets and TimeLapse-seq, Related to Figure 4.** (a) Functional enrichment analysis for common couplets. (b) Correlation between differential expression (x-axis) and differential binding sites (y-axis, top) and differential m6A sites (y-axis, bottom). (c) Distance between binding and modification for differential couplets for RM- and RBM15-specific couplets. (d) Schematic of TimeLapse-seq experiments. (e) PCA plot of “fraction new” for all s4U fed samples showing biological replicates cluster well together in terms of their kinetics, and the different conditions show differential clustering. (f) QC of raw T-to-C mutation rates for all samples sequenced, showing s4U-fed samples have significantly higher T-to-C mutation rates than the unfed samples, which represent background mutation rates. The data passed all internal QC tests performed by bakR. (g) Dot blot showing similar degrees of s4U incorporation in all sample types. (h) mRNA half-life for non-couplet genes in HEL-parental, HEL-RBM15, and HEL-RM cells (***p<0.001, Wilcoxon test). (i) *left panel:* Volcano plot of kdeg in HEL-RM vs HEL-RBM15 cells. Blue points have significantly higher degradation in HEL-RM than HEL-RBM15; orange points have significantly lower degradation rates in HEL-RM than HEL-RBM15 (FDR > 0.05, L2FC > 0). *right panel:* Volcano plot of ksyn in RM vs RBM15. Orange points have significantly higher synthesis in HEL-RM than HEL-RBM15. Blue points have significantly lower synthesis rates in HEL-RM than HEL-RBM15 (FDR > 0.05, L2FC > 0). Black points are RM couplet genes that had significant differential kdeg or ksyn in RM vs RBM15. (j,k) Mechanistic categorization of TimeLapse-seq data comparing HEL-RM to HEL-RBM15. Non-couplets are indicated in grey; couplets are represented in yellow (RBM15) and red (RM). (l) Half-lives of RM couplets compared to non-couplets in RM vs parental cells (p<0.001, Wilcoxon test). (m) RNA expression changes of RM, RBM15, and common couplets.

Figure S5

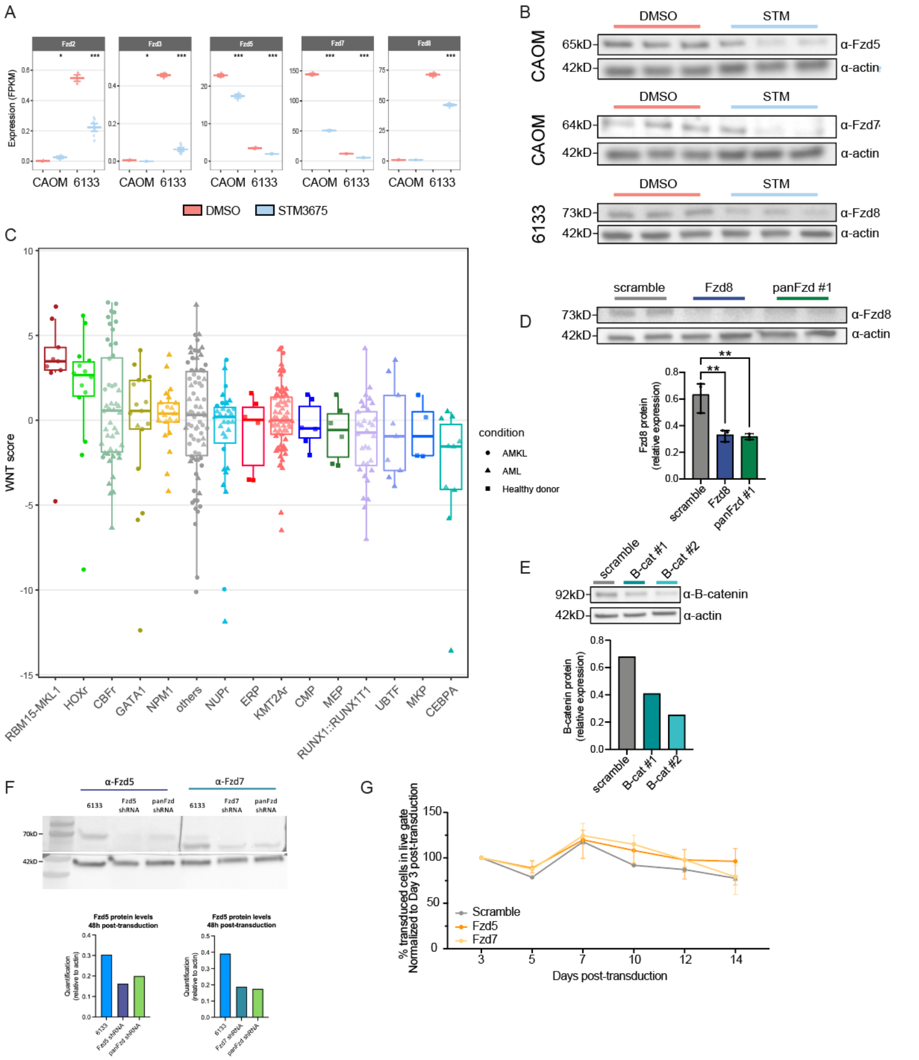


**Figure S5. Additional data on Wnt/Fzd expression and knockdown, Related to Figure 5.** (a) FPKM of most highly expressed Fzd genes in 6133 and CAOM cells with Vehicle or STM3675 treatment. Stars represent p-value after the Quasi-Likelihood F-Test. (* p<0.05, **p<0.01, ***p< 0.001). (b) Western blots showing decrease of Fzd proteins upon Mettl3 inhibition. (c) Wnt score across AMKL and AML genotypes as well as CMP, MEP, MKP, and ERP from healthy donors. Boxes represent median and quartiles, whiskers extend 1.5 times the IQR from the box edges. Western blots showing validation of shRNA-mediated knockdown of (d) Fzd8 (**$p<0.01$, one-way ANOVA with Bonferroni correction for multiple comparisons) and (e) β-catenin in 6133 cells. (f) Western blots showing validation of shRNA-mediated knockdown of Fzd5 and Fzd7 in 6133 cells. (g) 6133 cells transduced with lentivirus shRNA targeting Fzd5 and Fzd7 have no change in growth compared to scrambled control ($p>0.05$). Bars represent mean and range.
